# Effects of Spike Mutations in SARS-CoV-2 Variants of Concern on Human or Animal ACE2-Mediated Virus Entry and Neutralization

**DOI:** 10.1101/2021.08.25.457627

**Authors:** Yunjeong Kim, Natasha N Gaudreault, David A. Meekins, Krishani D Perera, Dashzeveg Bold, Jessie D. Trujillo, Igor Morozov, Chester D. McDowell, Kyeong-Ok Chang, Juergen A. Richt

## Abstract

SARS-CoV-2 is a zoonotic agent capable of infecting humans and a wide range of animal species. Over the duration of the pandemic, mutations in the SARS-CoV-2 Spike protein (S) have arisen in circulating viral populations, culminating in the spread of several variants of concern (VOC) with varying degrees of altered virulence, transmissibility, and neutralizing antibody escape. In this study, we employed lentivirus-based pseudotyped viruses that express specific SARS-CoV-2 S protein substitutions and cell lines that stably express ACE2 from nine different animal species to gain insights into the effects of VOC mutations on viral entry and antibody neutralization capability. All animal ACE2 receptors tested, except mink, support viral cell entry for pseudoviruses expressing the parental (prototype Wuhan-1) S at levels comparable to human ACE2. Most single S substitutions (e.g., 452R, 478K, 501Y) did not significantly change virus entry, although 614G and 484K resulted in a decreased efficiency in viral entry. Conversely, combinatorial VOC substitutions in the S protein were associated with significantly increased entry capacity of pseudotyped viruses compared to that of the parental Wuhan-1 pseudotyped virus. Similarly, infection studies using live ancestral (USA-WA1/2020), Alpha, and Beta SARS-CoV-2 viruses in hamsters revealed a higher replication potential for the Beta variant compared to the ancestral prototype virus. Moreover, neutralizing titers in sera from various animal species, including humans, were significantly reduced by single substitutions of 484K or 452R, double substitutions of 501Y-484K, 452R-484K and 452R-478K and the triple substitution of 501Y-484K-417N, suggesting that 484K and 452R are particularly important for evading neutralizing antibodies in human, cat, and rabbit sera. Cumulatively, this study reveals important insights into the host range of SARS-CoV-2 and the effect of recently emergent S protein substitutions on viral entry, virus replication and antibody-mediated viral neutralization.

**Author summary:** Cells stably expressing ACE2 from various animals and a lentivirus-based SARS-CoV-2 pseudotyped virus assay were established to study SARS-CoV-2 cell entry. The results demonstrated that ACE2 from a wide range of animal species facilitate S-mediated virus entry into cells, which is supported by *in silico* data as well as natural and experimental infection studies. Pseudotyped viruses containing mutations in the RBD of S representative of the Alpha, Gamma, and especially Beta, variants of concern demonstrated that certain mutations are associated with increased viral entry compared to the parental S. The Beta variant was also observed to have a replicative advantage *in vitro* and *in vivo* compared to the prototype virus. Pseudotyped viruses containing combinatorial substitutions of 501Y-484K-417K, 614G-501Y-484K and 614G-501Y-484K-417N increased viral entry via ACE2 across multiple species. The 501Y or 478K single substitution did not significantly affect neutralizing capacity of immune sera compared to the prototype strain, but the addition of 484K or 452R substitutions significantly reduced the neutralizing titers.

## Introduction

Severe acute respiratory syndrome coronavirus 2 (SARS-CoV-2), the etiological agent of COVID-19 disease, unexpectedly emerged in late 2019 and has spread throughout the world, infecting over 200 million people worldwide and causing over 4 million deaths as of August 2021 [1]. The zoonotic origin and intermediate hosts of SARS-CoV-2 are still unclear, although bats are considered a likely source based on numerous SARS-CoV-2-related bat coronaviruses found in Southeast Asia [2–4]. It is now increasingly apparent that SARS-CoV-2 has the capacity to infect several animal species besides humans, generating rising concerns that domestic and wild animals may become secondary reservoirs of the virus [5–7]. Outbreaks of COVID-19 in hundreds of mink farms in the EU [8], where identification of human-to-mink and mink-to-human virus transmissions [9, 10] as well as mink-associated variants led to the culling of over 20 million minks in Denmark, underscored the importance of identifying and assessing the risks associated with the new pandemic for animal and human health [11–14]. Other animal species, including cats, dogs, ferrets, hamsters, non-human primates, white-tailed deer, mice, cattle, pigs, tree shrews, rabbits, raccoon dogs, and fruit bats, have also been investigated for their susceptibility to SARS-CoV-2 infection [15]. Reports from natural and experimental infection studies determined a wide range of susceptibility in several domesticated (farm or companion) animals or wildlife to SARS-CoV-2 infection [8, 16–27].

SARS-CoV-2 is an enveloped, positive-sense RNA virus that belongs to the family *Coronaviridae.* RNA viruses are prone to high mutation rates, giving rise to new variants over time, although the mutation rate of coronaviruses is lower than many other RNA viruses due to proofreading activity of their replicative complex [28, 29]. Some virus variants possess notable changes in virus transmissibility, virulence, or other characteristics that are important in host defense, such as immune evasion. Since the emergence of COVID-19, multiple variants of SARS-CoV-2 have been identified and have largely replaced the prototype SARS-CoV-2 strain (Wuhan-Hu-1) [30, 31]. Currently, the World Health Organization designated Alpha (lineage B.1.1.7), Beta (B.1.351, B.1.351.2 and B.1.351.3), Gamma (P.1, P.1.1 and P.1.2) and Delta (B.1.617.2, AY.1 and AY.2) SARS-CoV-2 viruses as variants of concern (VOC) [20] as they are associated with increased risks to global public health. These variants contain multiple amino acid substitutions in the S protein, some of which have received special attention as they span the receptor binding domain (RBD) or the S1/S2 junction. Entry of SARS-CoV-2 to the target cells is mediated by the interaction of the S protein with its receptor, the ACE2 on the host cell membrane [3, 32, 33]. The RBD in the S protein is located on residues 319-541 and interacts with 25 conserved residues on human ACE2 [33, 34]. Cleavage of the S1/S2 junction (residues 613-705) of SARS-CoV-2 S protein by cellular proteases triggers fusion and viral entry into host cells [35, 36]. Due to its involvement in receptor binding, most neutralizing antibodies are directed against the RBD [37]. Mutations affecting the S protein, including the RBD, are of particular concern because they may enhance virus transmissibility and reduce antibody binding and immune protection, thus compromising vaccine efficacy [30]. In addition, the interaction between the cellular receptor and the virus, leading to virus entry into host cells, is one of the critical factors that determine host susceptibility to virus infection. With the recently emerging virus variants, it is critical to understand the impact and significance of such mutations on virus neutralization, which has wide-reaching implications on vaccine efficacy, and on animal susceptibility to SARS-CoV-2 to identify and manage risks of zoonotic/reverse zoonotic infections. Some of the key mutations found in SARS-CoV-2 VOC have been studied using pseudotyped viruses or recombinant viruses carrying mutant SARS-CoV-2 S proteins [38]; however, only limited information on the roles of the mutations for a broad range of animal species, as well as humans, is available so far.

Small animal models such as mice and Syrian Golden hamsters are available to study various aspects of SARS-CoV-2 infection [39]. Ancestral SARS-CoV-2 viruses can infect genetically engineered mice that express human ACE2, although unmodified mice are only permissive to mouse-adapted SARS-CoV-2 [40, 41] with the exception of SARS-CoV-2 variants containing the N501Y polymorphism in their S gene [42]. Hamsters are highly permissive to SARS-CoV-2 infection, and efficient virus replication and moderate to severe lung pathology are observed following virus replication, usually accompanied by weight loss and other clinical signs during acute infection [43–46]. Small animal models for COVID-19 have been used to study viral transmission, pathogenesis, immunity and for the evaluation of vaccines and therapeutic drugs and are also a suitable model for investigating virulence and infectivity of SARS-CoV-2 variants [47].

In this study we investigated the characteristics of key mutations found in Alpha, Beta, Gamma, or Delta variants (single or combinations of 614G, 501Y, 484K, 452R, 478K). Using lentivirus-based pseudotyped virus assays, the effects of key substitutions on virus entry into human and various animal-ACE2 expressing cells, and on the neutralizing activities of antisera from humans, cats, and rabbits were determined. Using the hamster model, infection studies were conducted to provide further understanding of the replication capacity of SARS-CoV-2 variants. Finally, structural models for the parental and mutant SARS-CoV-2 S protein RBD in complex with ACE2 from various animals were generated to probe the structural basis for host susceptibility and the effects of the mutations on the interactions between ACE2 and RBD. The presented results provide important insights into the impact of S protein mutations found in from emerging SARS-CoV-2 variants on cell entry in human and other animal species, and on virus replication and virus neutralization

## Results

### Entry of pseudotyped virus with SARS-CoV-2 S into HEK293T or CRFK cells expressing human or animal ACE2

Expression of ACE2 in HEK293T or CRFK cells that were stably transfected with a plasmid encoding the ACE2 protein from human and various animal species was confirmed using Western blot analysis (data not shown). Entry of pseudotyped viruses, measured by firefly luciferase, was comparable between HEK293T and CRFK cells expressing the same ACE2 construct. However, CRFK cells yielded more robust and consistent results than HEK293T; therefore, CRFK cells were subsequently used for pseudotyped virus entry assays. The results of the virus entry assays are shown in Figure 1A. Importantly, native CRFK cells that do not express exogenous ACE2, only inherent feline ACE2 (Mock), yielded negligible virus entry (Figure 1A), indicating that CRFK cells are suitable to determine the effects of exogenous heterologous ACE2 on viral entry. Expression of various animal ACE2 receptors in cells led to greatly enhanced entry of pseudotyped viruses expressing the parental SARS-CoV-2 S protein (Figure 1A), except for mink ACE2, which failed to show marked increase in virus entry compared to other ACE2s, with only a 31-fold increase over mock. Cellular entry of pseudotyped viruses in the presence of ACE2 receptors from various animal species ranged from an approximately 1,200-fold (horse/cat) to 3,000-fold (rabbit) increase in cellular entry compared to the mock control (no ACE2 transfection). Figure 1B shows a summary of the virus entry results using cells expressing different animal ACE2 receptors compared to that in cells expressing human ACE2. Virus entry levels for each ACE2 were considered high, medium, or low when > 80%, 10-80%, or 1-10% of virus entry in ACE2-expressing cells (compared to hACE2-expressing cells) was observed, respectively, based on the criteria suggested by Damas et al [18]. High levels of virus entry were observed in cells expressing ACE2 from human, dog, cow, hamster, or rabbit (Figure 1A and B), while medium levels of virus entry were seen in cells expressing ACE2 from cat, horse, camel, and white-tailed deer. Expression of mink ACE2 resulted in low virus entry. The overall trend of virus entry in cells expressing various animal ACE2 receptors was similar to the *in silico* predictions by Damas et al [18](Figure 1B).

**Figure 1.**
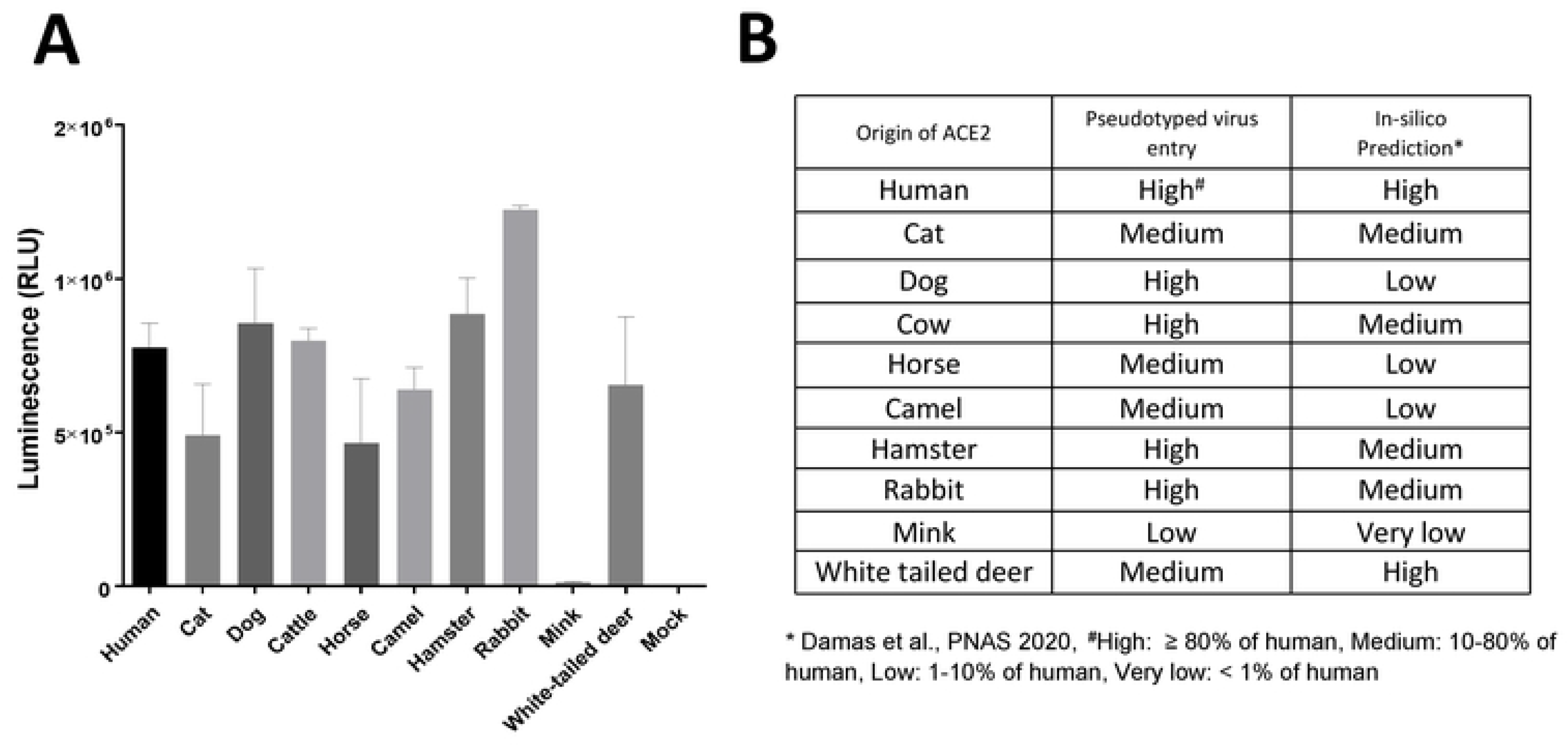
Effects of various ACE2 constructs on the entry of pseudotyped viruses carrying SARS-CoV-2 S into CRFK cells stably expressing ACE2 from various animal species. **A.** CRFK cells stably expressing various ACE2 receptors or mock cells (no ACE2 transfection) were infected with pseudotyped virus carrying the parental SARS-CoV-2 S protein. Following incubation of the cells with the pseudotyped virus for 48 h, cells were lysed, and luminescence units were measured. Each bar indicates the mean and the standard error of the means. **B.** Summary of the results from the pseudotyped virus entry assay in Figure 1A. Virus entry levels were considered high, medium, or low when > 80%, 10-80%, or 1-10% of virus entry in ACE2-expressing cells (compared to human ACE2 cells) was observed, respectively, based on the criteria suggested by Damas et al. [18]. \**In silico* predictions by Damas et al [18].

### Entry of pseudotyped virus expressing SARS-CoV-2 parental or mutant S in human ACE2-expressing CRFK cells

The pseudotyped virus preparations carrying single or multiple S mutations were quantitated and normalized by ELISA p24 lentivirus antigen measurement or by SARS-CoV-2 S protein expression for transduction of the cells. Virus entry of each pseudotyped virus carrying single or multiple substitutions of 417N, 452R, 478K, 484K, 501Y, or 614G on the S protein RBD was compared to that of parental pseudotyped viruses (no mutation in S gene) in cells expressing human ACE2. The single substitutions of 501Y, 452R, or 478K did not lead to a statistically significant difference in virus entry compared to parental virus (Figure 2A), except for 614G or 484K, which resulted in significantly reduced virus entry compared to that of the parental pseudotyped virus. Among the double substitutions (i.e., 614G-501Y, 501Y-484K, 452R-484K, or 452R-478K), only the 501Y-484K combination significantly increased pseudotyped virus entry compared to that of the parental pseudotyped virus. The addition of substitution 417N or 614G to the 501Y-484K combination, however, did not further increase the virus entry efficiency of pseudotyped virus compared to the 501Y-484K double substitution, unless both 417N and 614G were combined with 501Y-484K in a quadruple combination (417N-484K-501Y-614G). Interestingly, when 501Y was combined with 614G (614G-501Y double substitution), an increase of virus entry was observed, similar to the level of parental virus. Virus entry capacity was further enhanced by the addition of 484K (614G-501Y-484K) or 484K-417N (614G-501Y-484K-417N). Likewise, the combination of 501Y and 484K led to significantly increased virus entry compared to the parental virus, suggesting that the 501Y substitution is important in negating the suppressive effects of the 484K and 614G single mutations (Fig. 2A). The reduced virus entry due to the 484K substitution was also restored to the level of the parental virus entry when combined with the 452R substitution (Fig. 2A). However, the 452R-478K double mutation did not lead to enhanced virus entry compared to the 452R or 478K single mutations. In CRFK cells expressing no exogenous ACEs (native feline ACE2-expressing CRFK cells), a significant decrease or increase in pseudotyped virus entry was observed with the 614G single mutation or the 614G-501Y-484K-417N quadruple mutation, respectively (Fig. 2B). However, the overall magnitude of pseudotyped virus entry in non-transfected CRFK cells was very low, regardless of the presence or absence of S protein mutations, which confirms that non-transfected CRFK cells are poorly supportive of SARS-CoV-2 S-pseudotyped virus entry.

**Figure 2.**
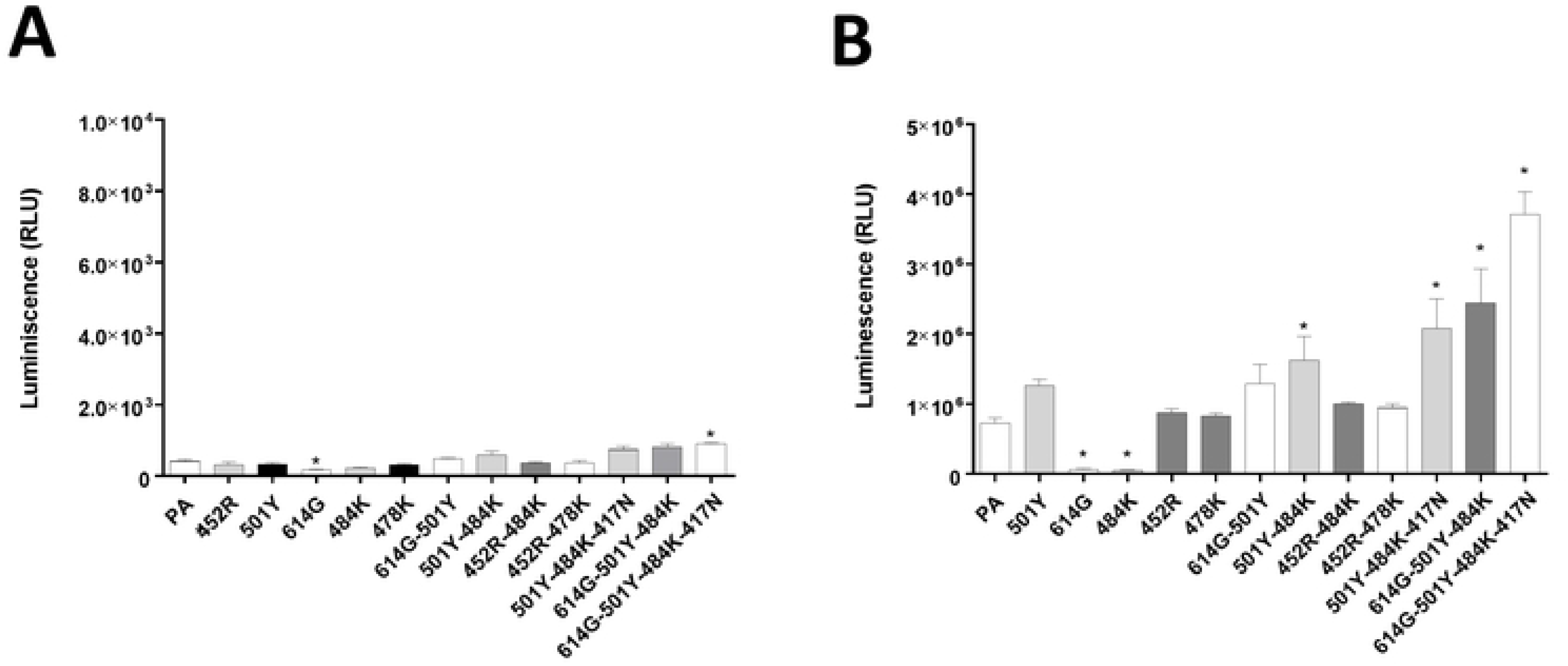
Entry of pseudotyped viruses carrying SARS-CoV-2 S with single or multiple substitutions on the RBD site of the S protein into non-transfected CRFK or CRFK cells stably expressing human ACE2. CRFK cells expressing human ACE2 (**A**) or mock (non-ACE2 transfected) CRFK cells (**B**) were infected with pseudotyped viruses with single or multiple substitutions. Following incubation of the cells for 48 h, luminescence units were measured. Each bar indicates the mean and the standard error of the means. PA indicates parental pseudotyped virus (no mutation in the S protein). One way ANOVA on the log10-transformed raw relative luminescence units were used to compare the parental (PA) and other groups. Statistical differences between mutation and the parental virus groups are indicated with * (p<0.05).

**Figure. 3.**
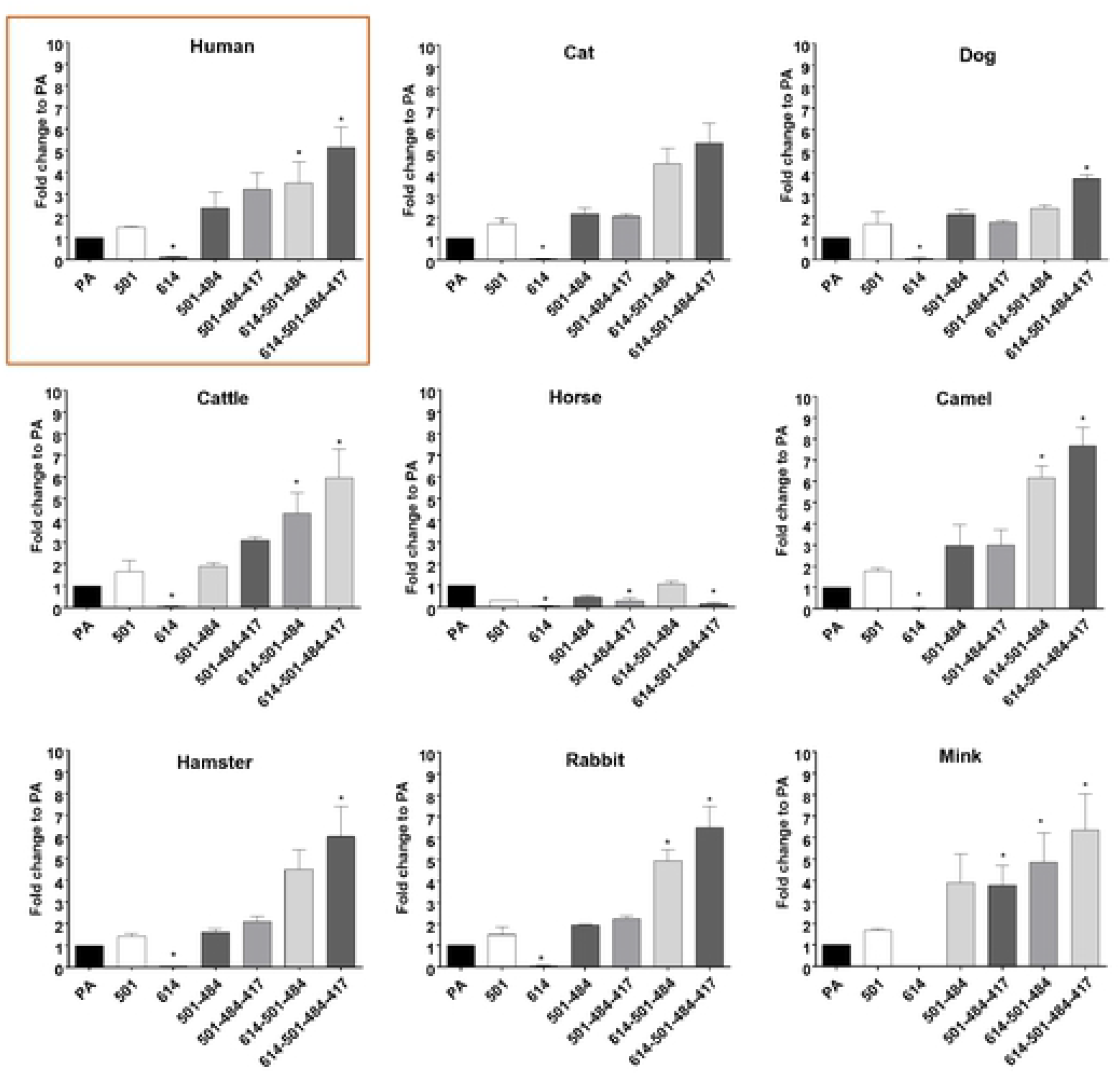
Entry of pseudotyped viruses carrying SARS-CoV-2 S with single or multiple mutations on the RBD site of S protein into CRFK cells expressing ACE2 of various species. CRFK cells expressing ACE2 were infected with pseudotyped viruses expressing single or multiple S protein substitutions. Following incubation of the cells for 48 h, cells were lysed, and relative luminescence units were measured. Each mutant pseudotyped virus was compared with the parental pseudotyped virus (PA), and data are presented as the fold-change to PA. One way ANOVA on the log10 transformed raw relative fluorescence units were used to compare the parental and other groups. Statistical differences between mutation and the parental virus groups are indicated with * (p<0.05).

### Entry of pseudotyped virus carrying SARS-CoV-2 parental or mutant S proteins in various ACE2-expressing CRFK cells

In this experiment, we compared the entry of pseudotyped viruses with parental or mutant S into cells expressing ACE2 from various animal species, including humans. Overall, the trend of change in virus entry among various pseudotyped viruses is similar in all tested cells expressing various animal ACE2 receptors. The quadruple 614G-501Y-484K-417N substitution showed the highest fold-change compared to the parental S (no mutation), followed by the triple combination 614G-501Y-484K. The 501Y-484K and 501Y-484K-417N substitutions led to a moderately increased virus entry compared to the parental S, but without a statistically significant difference. The 614G single mutation led to a decrease in virus entry in cells expressing human (see also Figure 2A) and animal ACE2. Notably, even in mink ACE2-expressing cells, which support limited virus entry compared to other ACE2s, a similar trend was observed with pseudotyped viruses with single and multiple substitutions. Interestingly, relatively little change was observed in virus entry among parental and mutant pseudotyped viruses in horse ACE2-expressing cells. These results suggest that the effects of these mutations in the RBD region of the S protein for virus entry are shared among a wide range of animal ACE2 receptors.

### Replication and infection dynamics of SARS-CoV-2 variant strains

To gain insights into the consequences of VOC mutations, replication kinetics of parental lineage A (WA1/2020), B.1, B.1.1.7 (Alpha), and B.1.351 (Beta) SARS-CoV-2 strains in CRFK-human ACE2 or CRFK-hamster ACE2 cells were determined (Fig. 4 A, B). Although a 0.01 MOI was intended, back-titration of the inoculum, represented as time 0, showed the B.1.351 strain at a significantly lower input titer than the lineage A parental prototype strain. However, titers for the lineage A virus were lower at 24, 48 and 72 hpi compared to the B.1, B.1.1.7 and B.1.351 strains for both CRFK-human ACE2 or CRFK-hamster ACE2 cells. Moreover, the B.1.351 titers were higher than the parental lineage A virus in CRFK-human ACE2 cells at 24, 48 and 72 hpi, and B.1.351 was the dominant virus strain in the CRFK-hamster ACE2 cells at 48 and 72 hpi. In both cell lines, the B.1 strain dominated at 24 hpi. The B.1, B.1.1.7 and B.1.351 strains had relatively similar titers in the human ACE2 cells at 48 and 72 hpi, while more variation was observed between these strains at the same time points in CRFK-hamster ACE2 cells. All strains peaked at around 48 hpi in both cell lines, although generally lower titers were observed at all time points for CRFK-hamster ACE2 cells compared to CRFK-human ACE2 cells. Taken together, these results show that the variant viruses have a replication advantage over the parental lineage A strain in these cells.

**Figure 4.**
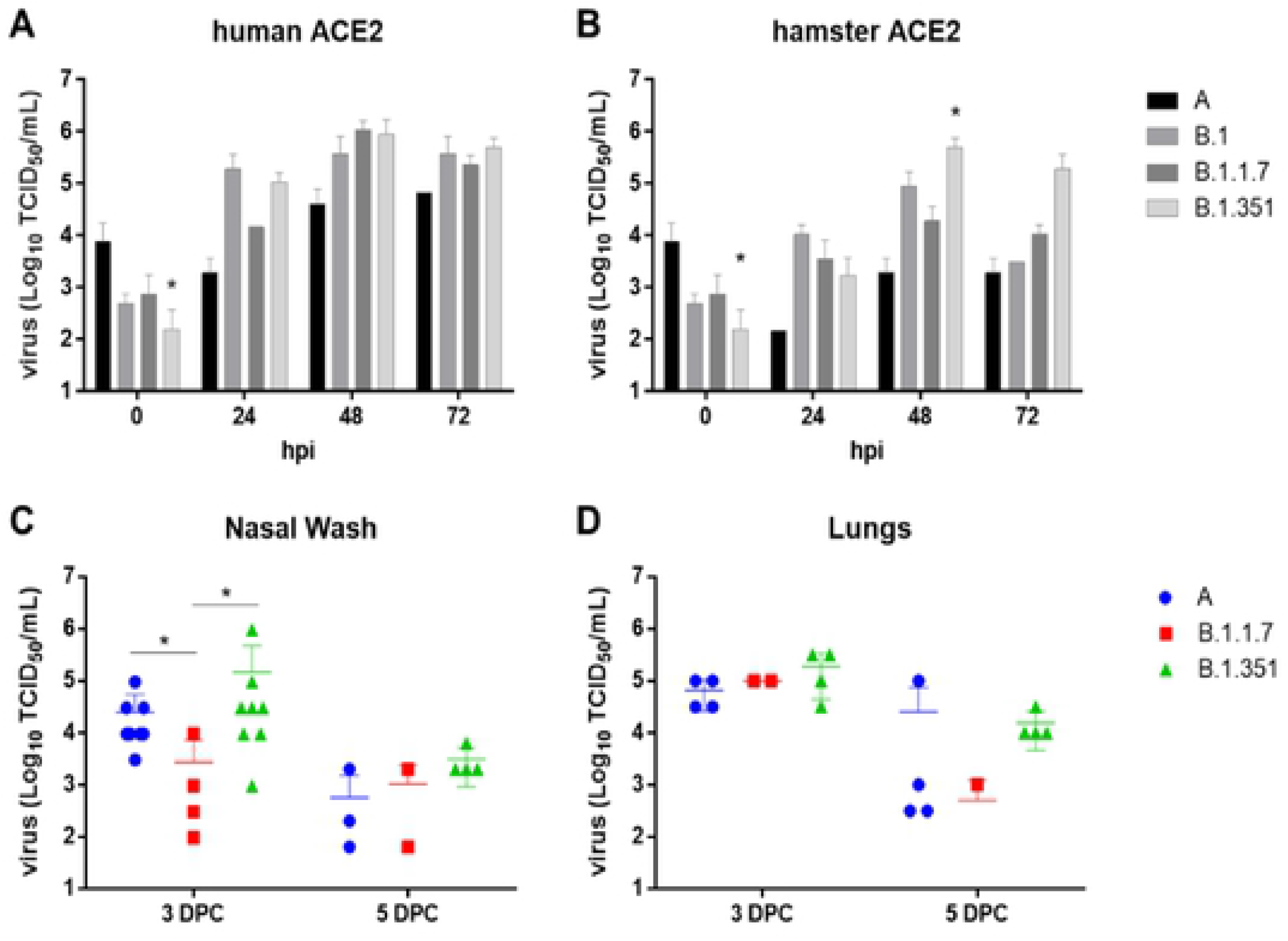
Replication and infection dynamics of SARS-CoV-2 variant strains in cells and hamsters. Replication kinetics were performed in CRFK-human ACE2 (A) or CRFK-hamster ACE2 (B) cells. Virus titrations were performed on infected cell culture supernatants collected at 24, 48, and 72 hours post infection (hpi), and the virus inoculum (timepoint 0). Virus titrations were also performed on the nasal wash (C) and lungs (D) collected from hamsters inoculated intranasally with 1×10^5^ TCID_50_ of each virus strain at 3- and 5-days post challenge (DPC). Mean and standard deviations are shown for each data set. Two-way ANOVA of transformed virus titer data was performed, and statistical differences of the variants compared between each of the strains per time point are indicated by * (p<0.05).

Replication ability of the parental lineage A, Alpha VOC B.1.1.7 and Beta VOC B.1.351 strains were analyzed in hamsters inoculated intranasally with 1×10^5^ TCID_50_ of the respective viruses (Fig. 4 C,D). Mean virus titers of nasal washes collected from the lineage A- and Beta VOC B.1.351-infected groups were significantly higher than the B.1.1.7 infected group at 3 DPC, but not significantly different at 5 DPC. Mean virus titers of lung homogenates were similar for all groups at 3 and 5 DPC although the B.1.351 infected individuals had generally higher nasal wash and lung titers compared to the linage A- and B.1.1.7-infected animals. Together these data support the notion that the Beta VOC B.1.351 strain has a replicative advantage in hamsters compared to the parental lineage A and B.1.1.7 strains.

### Neutralizing activities of convalescent human sera or post-infection/vaccination sera from cat, and rabbit against pseudotyped viruses with single S protein mutations

The effects of single mutations on the S protein in pseudotyped viruses in the neutralizing activity of various sera from human, cat, and rabbit were assessed using human ACE2-expressing cells. The results are shown in Figure 5 and Table 1. The negative control sera from human, cat and rabbit did not neutralize any of the pseudotyped viruses (starting at a 12.5 dilution), as expected (Figure 5C, F and I). The 501Y or 478K single substitutions did not affect the neutralizing activity of all tested sera (Figure 4A-I, Table 1). However, significantly reduced neutralizing activities of human and cat sera were observed against pseudotyped viruses carrying 484K or 452R single substitutions. In contrast, the reduction in neutralization due to the 484K or 452R single mutations was not apparent in the rabbit sera, except for the rabbit 7A serum against 452R.

**Figure 5.**
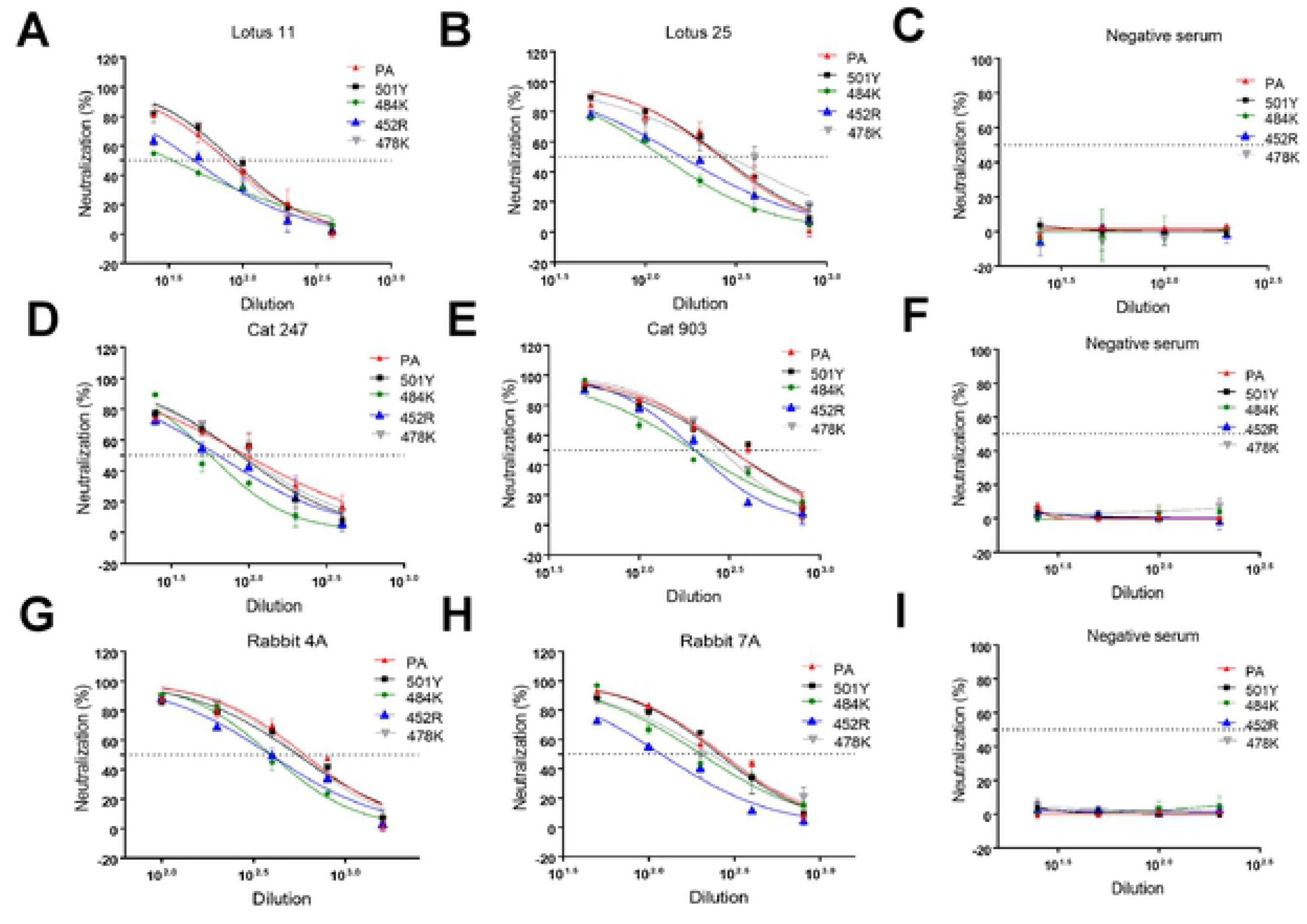
Neutralizing activity of various human, cat, and rabbit (sera against pseudotyped viruses carrying SARS-CoV-2 parental or single mutant S in cells expressing human ACE2. CRFK cells expressing human ACE2 were infected with pseudotyped viruses and tested for neutralizing activity against various human (A, B, C), cat (D, E, F), and rabbit (G, H, I) sera. Following incubation of the cells for 48 h, cells were lysed, and the relative luminescence units were measured. Each bar indicates the mean and the standard error of the means. PA indicates parental pseudotyped virus.

**Table 1.** Neutralizing antibody titers of various human, cat, and rabbit sera against pseudotyped viruses carrying SARS-CoV-2 parental (PA) or mutant S proteins tested with human ACE2-expressing cells. One-way ANOVA on the neutralizing titers was used to compare the parental and mutant groups. Statistical differences between the mutations and the parental group are indicated with * (p<0.05).

|  | Humans |  |  | Cats |  |  | Rabbits |  |  |
| --- | --- | --- | --- | --- | --- | --- | --- | --- | --- |
|  | Lotus 11 | Lotus 25 | Neg | #247 | #903 | Neg | #4A | #7A | Neg |
| **PA** | 78.53±2.3 | 275.65±6.3 | < 12.5 | 96.53±4.2 | 328.60±21.1 | < 12.5 | 534.05±77.6 | 270.10±16.7 | < 12.5 |
| **501Y** | 85.71±5.3 | 271.05±34.3 | < 12.5 | 89.32±5.9 | 322.05±30.1 | < 12.5 | 550.95±16.4 | 259.05±22.6 | < 12.5 |
| **484K** | 33.56±0.5* | 125.00±4.3* | < 12.5 | 56.82±4.3* | 201.40±2.3* | < 12.5 | 395.30±22.7 | 201.40±2.3 | < 12.5 |
| **452R** | 47.11±5.3* | 161.25±12.7* | < 12.5 | 64.18±6.9* | 204.15±4.1* | < 12.5 | 391.65±17.7 | 117.75±0.8* | < 12.5 |
| **478K** | 78.73±0.3 | 290.85±14.9 | < 12.5 | 92.37±3.4 | 282.35±5.8 | < 12.5 | 563.40±14.5 | 239.55±55.0 | < 12.5 |
| **501Y-484K** | 34.38±2.3* | 84.98±2.3* | < 12.5 | 29.16±0.6* | 135.60±16.7* | < 12.5 | 265.10±30.7* | 95.06±10.4* | < 12.5 |
| **452R-484K** | 25.62±4.0* | 114.02±23.2* | < 12.5 | 37.19±0.6* | 143.45±9.4* | < 12.5 | 201.85±9.2* | 111.10±3.0* | < 12.5 |
| **452R-478K** | 27.40±0.6* | 89.57±5.2* | < 12.5 | 39.634±7.9* | 103.34±9.9* | < 12.5 | 228.20±12.9* | 97.63±10.5* | < 12.5 |
| **501Y-484K-417N** | 25.9±0.0* | 90.22±1.6* | < 12.5 | 31.98±2.0* | 103.20±3.0* | < 12.5 | 197.05±3.5* | 110.50±4.9* | < 12.5 |

### Neutralizing activities of convalescent or post-infection sera from human, cat, and rabbit against pseudotyped viruses with double or triple S protein mutations

The results of the virus neutralization assays (VNAs) are summarized in Figure 6 and Table 1. Interestingly, when the 501Y or 478K mutation, which did not affect neutralization as a single substitution (Figure 5), is combined with 484K or 452R (i.e., 501Y-484K, 452R-478K), neutralizing activities of all tested sera were significantly decreased when compared to the parental pseudotyped virus. Moreover, the double substitution 452R-484K and the triple substitution 501Y-484K-417N also resulted in significant reduction of neutralizing activities of all tested sera compared to the parental pseudotyped virus. Each of these double and triple substitutions showed comparable neutralizing titers in all tested sera. These results from single and multiple substitutions in the S protein suggest that positions 484K and 452R are particularly important for evading neutralizing antibodies in human, cat, and rabbit sera.

**Figure 6.**
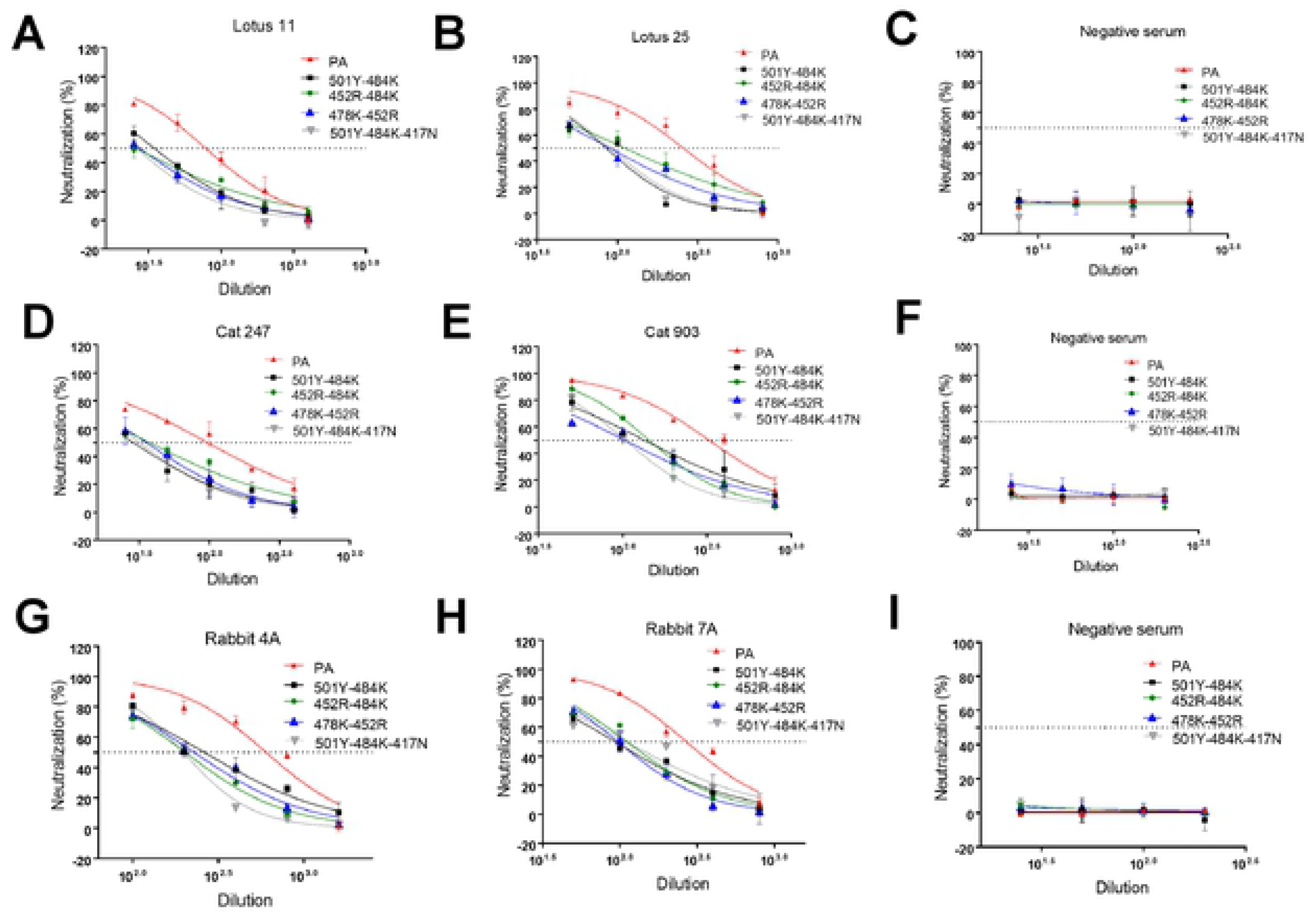
Neutralizing activity of various human, cat, and rabbit sera against pseudotyped viruses carrying SARS-CoV-2 parental (PA) or multiple S mutations in cells expressing human ACE2. CRFK cells expressing human ACE2 were infected with pseudotyped viruses and tested for neutralizing activity against various human (A, B, C), cat (D, E, F), and rabbit (G, H, I) sera. Following incubation of the cells for 48 h, cells were lysed, and the relative fluorescence units were measured. Each bar indicates the mean and the standard error of the means. PA indicates parental pseudotyped virus.

### Structural modeling insights into the ACE2-RBD interaction of different species

Structural modelling (Figure 7A) showed that parental (PA) SARS-CoV-2 RBD forms hydrogen bonding interactions (dotted lines) at the following RBD positions: (i) N501 with human (h)ACE2-Y41 and hACE2-K353; (ii) K417 with hACE2-D30; and (iii) E484 with hACE2-K31 (Figure 7B). In the triple 501Y-484K-417N mutant, Y501 interacts with hACE2-Y41 and hACE2-K353 and the salt bridges between N417 and K484 and the ACE2 receptor are lost (Figure 7B). No significant structural differences in the RBD-ACE2 interactions were observed for the PA or mutated RBD for ACE2 from cat, cattle, hamster, rabbit, or white-tailed deer (data not shown). However, dog and mink ACE2 receptors contain an ACE2-H34Y substitution (Figure 7C) that may increase binding affinity in the presence of the S Protein K417N substitution due to potential interactions between the hydroxyl group of tyrosine and the amino group of asparagine (Figure 4B). Moreover, horse ACE2 contains a Y41H substitution (Figure 4C) that does not provide the interactions necessary for strong binding of either the PA N501 or mutant Y501 residues (Figure 4B). Lastly, camel ACE2 contains a K31E substitution (Figure 4C), which may provide a basis for polar contacts in the region in the presence of the S protein K484 substitution. Overall, analysis of these structural models provides insights into the cell entry of the parental and mutated S proteins in the presence of ACE2 receptors from the different species.

**Figure 7.**
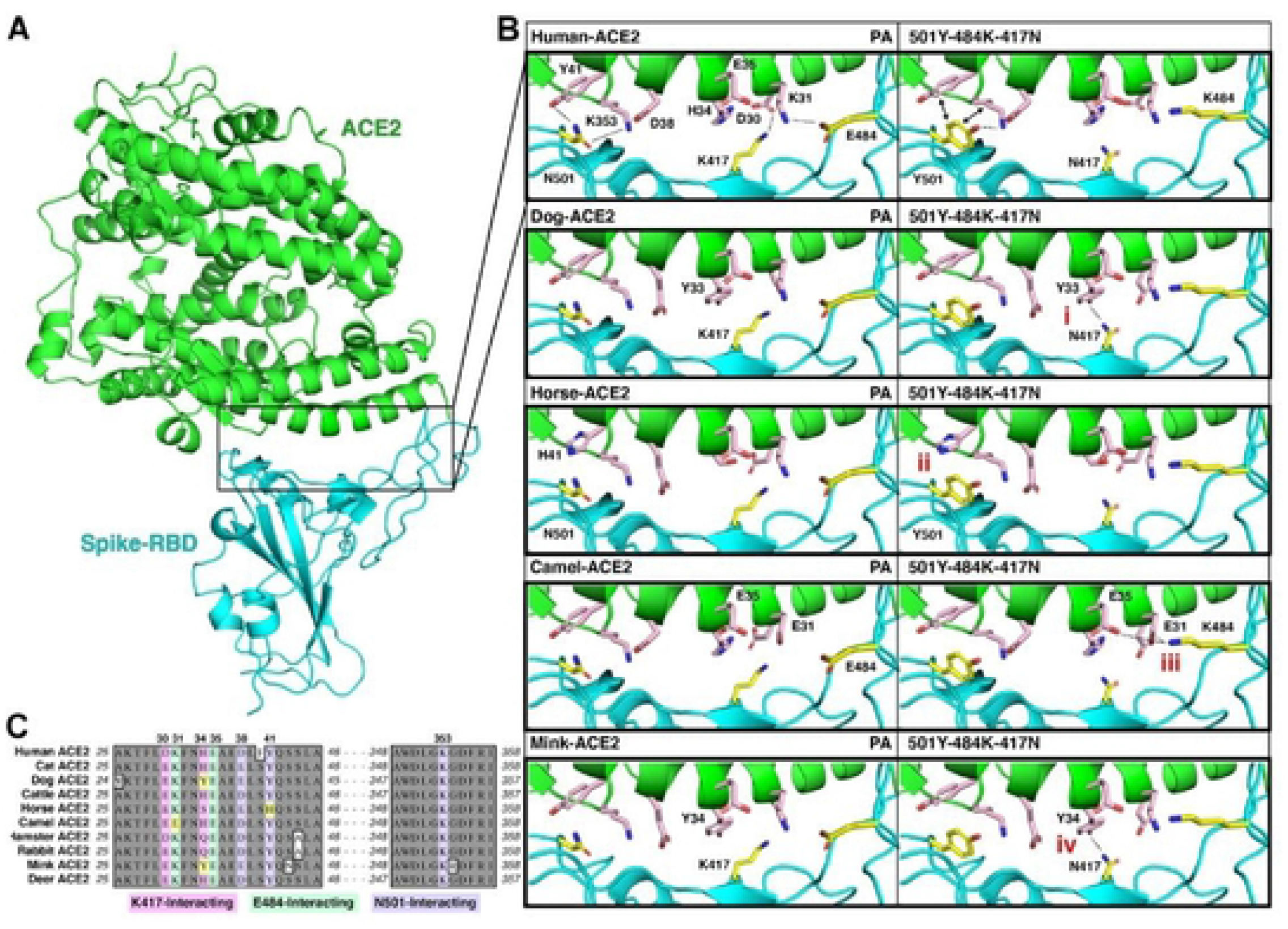
Structural modeling insights into the ACE2-RBD interaction in different species. **A.** Illustration of the human ACE2 (green) interaction with the SARS-CoV-2 RBD (cyan) as determined by Protein DataBank (PDB) entry 6GLZ [91]. The inset outlines the location of the ACE2-RBD interface illustrated in (B). **B**. Summary of significant differences from the human ACE2-RBD interface in the investigated species, highlighting notable differences in (i) dog, (ii) horse, (iii) camel, and (iv) mink. **C**. Alignment of the ACE2 amino acid sequences of each species (Supplemental Table 1), with K417 (pink), E484 (green), and N501 (purple) interacting residues highlighted. Significant differences outlined in (B) are highlighted in yellow.

## Discussion

Since the unexpected emergence of SARS-CoV-2 in human populations, extensive efforts have been directed towards both elucidating the risks associated with emerging virus variants and identifying susceptible animal species to better understand the zoonotic/reverse zoonotic implications of the pandemic. In our study, we used pseudotyped virus assays to elucidate the roles of ACE2 from various animal species including humans, in viral entry, which is a central event determining host susceptibility to SARS-CoV-2 infection. Using the S protein from the ancestral prototype (parental) SARS-CoV-2 stain (Wuhan-Hu-1), we found that several animal ACE2 receptors can efficiently interact with the SARS-CoV-2 S protein to allow virus entry into cells. The efficiency of virus entry among animal ACE2 receptors tested are not remarkably different from that of human ACE2, except for mink ACE2 which was consistently associated with comparatively low virus entry. Many animal species have been reported to be susceptible to SARS-CoV-2 infection either in experimental studies or natural infection as evidenced by clinical disease, viral replication in the respiratory tract and other organs, viral shedding/transmission, or seroconversion; these include domestic and large captive cats, dogs, cattle, mink, ferrets, otters, fruit bats, non-human primates, New Zealand white rabbits, hamsters, deer mice, bushy-tailed woodrats, striped skunks, and white-tailed deer [48–52]. Other animal species either have not been tested or showed no consistent evidence of active viral infection. Among them, cats and dogs have been of particular interest due to their proximity to humans. These companion animals can be infected by SARS-CoV-2 in natural and experimental settings, and usually remain asymptomatic although some develop mild respiratory disease [23, 53–55]. Overall, our pseudotyped virus entry results are consistent with previous animal susceptibility studies with all animal ACE2 receptors (human, cat, dog, cow, horse, camel, hamster, rabbit, mink, and white-tailed deer) tested in this report allowing virus entry [48–52]. Although there are currently no reports of natural or experimental infection in horses and camels, there have been concerns that SARS-CoV-2 may infect these animals based on predictions from structural *in silico* analyses or cell-to-cell fusion assays using pseudotyped virus [16–18]. Our results regarding the horse and camel ACE2 receptors and pseudotyped viruses provide a further impetus to study viral susceptibility in these animal species; however, our structural modeling, coupled with previous experimental evidence [56], indicates that the horse ACE2 Y41H substitution may confer resistance to RBD binding of both parental and mutated S proteins. An experimental infection study of cattle revealed that SARS-CoV-2 infection in this species may occur but does not appear to be robust, which seems to support the results of pseudotyped virus assays conducted by us and others [16, 57]. Interestingly, mink ACE2 was predicted to have a weak interaction with S protein in a previous *in silico* analysis study [18]; similarly, our pseudotyped virus entry assay showed that mink ACE2 allowed viral entry, although at a relatively lower level compared to other animals or human ACE2. This is somewhat surprising because mink are highly susceptible to SARS-CoV-2 infection, leading to a significant number of outbreaks of COVID-19 in mink farms with high morbidity/mortality [8, 19]. It is likely that an unknown disparity exists between virus entry mediated by pseudotyped viruses and native cell-virus interaction for mink. Structural models were generated to gain insight into the interaction between the S protein and selected animal ACE2 with a focus on the residues interacting with K417, E484 and N501 on the S protein (Fig. 7). The respective ACE2 residues are mostly conserved with minor variations among human and animal ACE2s, which is line with the pseudotyped virus assay results obtained in this study.

We also examined the effects of various mutations (417, 452, 478, 484 and 501) in the RBD, found in the Alpha (614G-501Y), Beta (614G-501Y-484K-417N), or Delta variants (452R-478K or 452R-484K), on entry of cells expressing human or animal ACE2 receptors using pseudotyped viruses. SARS-CoV-2 variants carrying 614G have replaced the prototype 614D virus and now are part of all major variants [58, 59], most likely because 614G is associated with enhanced fitness in susceptible cell lines including human airway cell culture [59, 60]. Th 614G virus was also shown to enhance replication in the upper respiratory tract and transmission in infected hamsters [60, 61], although this was not observed in hACE2 mice [60]. In human ACE2-expressing 293T cells, pseudotyped viruses carrying 614G alone have been reported to either increase [59, 62–65] or cause no change [38] in viral cell entry. In contrast to previous findings showing an increase in 614G cell entry in cells expressing human, cat or dog ACE2 orthologs [59], pseudoviruses carrying the 614G mutation alone consistently showed decreased cell entry across all species in our assays. Structural studies have indicated that 614G does not result in a higher affinity towards ACE2, but instead results in allosteric changes conducive towards a more open conformation of the RBD in which it is better positioned to interact with the ACE2 receptor [59]. The entry efficiency of the 484K single mutation alone has not yet been well studied. In our study using human ACE2 expressing cells, entry of the 614G or 484K mutant pseudotyped viruses was significantly decreased compared to the parental virus. In contrast, the 614G-501Y-484K (found in the Beta VOC) and 614G-501Y-484K-417N (found in Beta and Gamma VOC) mutations in the S protein increased virus entry compared to the parental pseudotyped virus. In a previous report [57], pseudoviruses with these mutations did not change virus entry in cells expressing human and various animal ACE2 receptors, with the exception of murine ACE2-expressing cells [57]. This observed difference in virus entry may be due to the different assay system including cell types, variance of assays, and other factors.

Our replication kinetics study using parental (lineage A), B.1, Alpha (B.1.1.7) and Beta (B.1.351) SARS-CoV-2 variants in CRFK cells stably expressing human ACE2 or hamster ACE2 revealed a comparatively higher replicative capacity of the Beta variant. This is consistent with the report of increased replication efficiency of Beta variant over Alpha variant in Vero cells [66]. In our experimental infection study using hamsters infected with parental virus (lineage A), Alpha (B.1.1.7) and Beta (B.1.351) VOC, more robust viral replication was observed in the nasal washes with the Beta VOC than the prototype virus or the Alpha VOC at 3 DPC, although the Beta variant inoculum titer was significantly lower than the inoculum dose of the other two viruses. This suggests that the Beta variant has a higher replication efficiency in susceptible nasal epithelial cells during the early phase of infection, which would be conducive towards increased virus shedding and transmission. The Alpha VOC appears to be transmitted highly efficiently [67, 68] and the Beta VOC has spread rapidly worldwide, which suggests that the Beta variant seem to have a replicative advantage over other variants previously circulating, although more data is required to confirm this [69].

The emerging VOC have been associated with an adverse impact on antibody neutralization capacity by convalescent and immunized humans, which may affect vaccine- and infection-induced protection, risks of reinfection, and the potency of therapeutic mAbs [70, 71]. The full set of signature mutations on and around the RBD of the S protein, as well as mutations in other genomic regions of SARS-CoV-2 VOC, contribute to the characteristics of each variant, and particular attention has been directed towards the roles of amino acid residues located on the S protein’s RBD, due to its direct interaction with ACE2. To understand the effects of key mutations of the Alpha, Beta, and Delta VOC, we tested respective pseudotyped viruses in virus neutralization assays with convalescent human and cat (from a challenge study with the SARS-CoV-2 USA-WA1/2020 strain [72]) sera, and hyperimmune rabbit sera (immunized with baculovirus-expressed prototype parental S protein). Substitution N501Y is found in Alpha, Beta and Gamma VOC and reported to have an increased binding efficacy to human ACE2 [73]. Our result showed the 501Y substitution did not significantly affect neutralization ability of all tested sera compared to the parental virus (PA); this is consistent with a previous report whereby recombinant SARS-CoV-2 virus (within the genetic background of USA WA1/2020) carrying the 501Y or 501Y-614G mutations did not affect neutralization ability of BNT162b6 vaccine-induced antibodies in human sera [74]. The 484K mutation has emerged independently in multiple variants and confers resistance to some mAbs targeting S protein receptor binding motif region [75, 76]. Adding the 484K mutation into pseudotyped viruses carrying the full set of the Alpha variant-associated mutations on the S protein led to a considerable loss in neutralizing activity of antibodies in BNT162b6-elicited or convalescent human sera [77]; this confirms our results using pseudotyped viruses carrying the 484K or 484K-501Y mutations. Likewise, combined S protein substitutions, such as 484K-614G, 484K-501Y-614G or 417N-484K-501Y-614G, reduced the neutralization ability of BNT162b6 vaccine-induced human sera and several therapeutic mAbs to recombinant virus in the genetic background of USA WA1/2020 [74]; similarly, it affected the neutralization ability of convalescent human sera to pseudotyped viruses carrying 417N-484K-501Y-614G. The substitutions 452R and 478K are found in the Delta variant and other variants under monitoring, such as the Epsilon variant, and have been on the rise in the US and Mexico since early 2021 [78]. Due to its relatively recent appearance, the impact of the 452R and 478K substitutions on the S protein have not yet been well studied; they have been predicted to negatively affect human ACE2 binding and shown to reduce neutralizing activity of antibodies from convalescent or vaccinated individuals [62, 79, 80]. In our pseudovirus assay, we observed significant reduction of neutralization activity of the tested sera with the pseudovirus carrying 452R, but not 478K. The negative effect of 452R on neutralization is also seen with the 452R-478K mutation, which significantly reduced neutralization activity of the sera. Interestingly, the neutralizing activity of sera with 452R-484K, 452-478, 501Y-484K and 501Y-484K-417N pseudoviruses were comparably decreased in our assays. The negative impact of the 501Y-484K-417N-carrying pseudovirus on neutralizing antibodies elicited with mRNA-based vaccines was also observed by others [81].

Our structural modelling showed potential changes in the interactions of RBD amino acid substitutions (501Y-484K-417N) and human ACE2, which may affect RBD binding. The structural modelling performed in this study is consistent with previous reports, indicating that the N501Y substitution increases the affinity for human ACE2 via the formation of new contacts with ACE2 residues Y41 and K353, and that the salt bridges between E484 and ACE2 K31, as well as K417 and D30 are lost upon substitution to 484K and 417N [82–84]. While many of the animal ACE2 sequences analyzed in this study (cat, cattle, hamster, and rabbit) contain ACE2-RBD interaction sites consistent with humans, there are amino acid differences in the ACE2 receptors of dog, mink, horse, and camel that could have significant effects on RBD-ACE2 interaction, especially when subjected to VOC. The H34Y ACE2 substitution found in dogs and mink was previously predicted to decrease RBD interaction, although several additional species with known susceptibility carry this substitution (e.g., cougar, lion, tiger) [85, 86]. Our modeling analysis suggests that the dog/mink Y34 residue in ACE2 could potentially increase RBD-ACE2 affinity when the RBD K417N substitution is present; however, the K417N substitution did not significantly increase pseudovirus entry in cells expressing ACE2 from dogs and mink. This inconsistency may limit the significance of the H34 residue of ACE2 and highlights the importance of wet lab research to supplement *in silico* predictions. Horse ACE2 contains a Y41H substitution, an amino acid which was shown by prior mutagenesis studies to abolish RBD binding [56, 87]. This contrasts with our results indicating that horse ACE2 with 41H can facilitate viral entry, albeit at a lower level. Notably, variant-associated S proteins did not increase cellular entry mediated by horse ACE2, indicating an overall deficiency for S binding for this species. Lastly, the camel ACE2 contains a K31E substitution that was previously shown to abolish ACE2 binding [87]; however, we observed robust viral entry in cells expressing the camel ACE2 receptor. Moreover, our structural predictions indicate that the K31E substitution in camel ACE2 may provide a basis for hydrogen bonding with 484K in the S variant. This is consistent with an increase in cell entry observed upon the E484K substitution, indicating that variants containing 484K may have increased susceptibility in camels. Overall, these structural insights provide some basis for understanding the ramifications of SARS-CoV-2 S mutations in emerging variants, although further structural and functional studies are required to confirm their effects in various animal species.

In summary, our results obtained from lentivirus-based system and hamster infection study showed that a wide range of animal ACE2 support pseudotyped virus entry, and the key mutations found in the VOC affect pseudotyped virus entry in cells expressing human or animal ACE2 as well as neutralizing sera of human, cat, and rabbit. The hamster infection study confirmed the replicative advantage of beta variant over the parental and alpha variant. The findings of this study highlight the importance of elucidating the roles of mutations and monitoring for evolving SARS-CoV-2 variants to assess their public health implications.

## Materials and Methods

### Animal care and ethics statement

All animal experiments were conducted in animal biosafety level 3 (BSL3) facilities at the Biosecurity Research Institute at Kansas State University according to protocols approved by the Institutional Animal Care and Use Committee at Kansas State University and the guidelines set by the Association for the Assessment and Accreditation of Laboratory Animal Care and the U.S. Department of Agriculture.

### Cells and plasmids

Human embryonic kidney 293 (HEK293), the Crandell-Rees feline kidney (CRFK) and Calu-3 cells were purchased from American Type Culture Collection (ATCC; Manassas, VA). Vero E6 cells expressing human TMPRSS2 (Vero-TMPRSS2) cells were obtained from Creative Biogene (Shirley, NY)[88]. Cells were maintained with either Dulbecco’s Modified Eagle Media (DMEM) or Eagle’s Minimal Essential Medium (MEM) supplemented with 5% fetal bovine serum, 100 U/ml penicillin and 100 μg/ml streptomycin, respectively. The codon-optimized cDNAs of the open reading frame of the human or animal ACE2 gene with FLAG tag were synthesized by Integrated DNA Technologies (Coralville, IA) and cloned into pIRES-Neo3 (Takara Bio, Mountain View, CA). For the ACE2 gene of white-tailed deer, because only partial ORF is available, the full ORF was constructed with human ACE2 gene. These plasmids were then designated as pIRES-Neo-(species) ACE2-FLAG. The animal species from which ACE2 gene sequences (listed in Supplemental Table 1) were derived are cat, dog, Arabian camel, European mink, horse, rabbit, cattle, Syrian golden hamster, and white-tailed deer. Pseudotyped viruses expressing SARS-CoV-2 S protein were generated by synthesizing the S gene which was truncated by 26 amino acids at the C-terminus, fused with HA tag by Integrated DNA Technologies, and cloned into plasmid pAbVec1 (Addgene, Watertown, MA), which were designated as pAbVec-SARS2-S. The parental S gene sequence was the prototype SARS-CoV-2 S gene from Wuhan (GenBank ID: YP_009724390.1). This clone was then used to generate single or multiple mutations in the RBD of S gene with site-directed mutagenesis kit (Agilent, Santa Clara, CA) using primers listed in the supplemental Table 2 and designated as pAbVec-SARS2-S (mutant). Single mutations in the RBD include N501Y (alpha variants), E484K, K417N, T478K and L452R, and multiple mutations include N501Y+E484K (gamma variants), L452R+E484K (delta variants), L478K+L452R (delta variants), N501Y+E484K+ K417N (beta variants), and D614G+N501Y-E484K+K417N (beta variants). Each mutation was confirmed by Sanger sequencing analysis.

### Anti-SARS-CoV-2 antibodies from humans, cats, and rabbits

Convalescent sera (Lotus 11 and 25) from COVID-19 patients were obtained from Dr. Thomas Rogers from the Scripps Research Institute, San Diego, CA, USA. Cat sera (Cat 247 and 903) were collected from cats enrolled in SARS-CoV-2 re-infection studies [89]. Hyperimmune rabbit sera (Rabbit 4A and 7A) were obtained by immunizing rabbits with SARS-CoV-2 recombinant S baculovirus-expressed proteins based on the prototype Wuhan isolate. Negative sera from each species were also included in the study.

### Generation of CRFK cells stably expressing human or animal ACE2

CRFK cells, plated the previous day, were transfected with pIRES-Neo-human (or cat, dog, cattle, horse, camel, hamster, rabbit, mink, white-tailed deer) ACE2-FLAG. The transfected cells were then subsequently selected in the presence of 1 mg/ml G418. Expression of each species’ ACE2 receptor in the cells was confirmed by Western Blot analysis using antibody to human ACE2 (Abcam, Waltham, MA). Parental CRFK cells served as a control (Mock).

### Generation of SARS-CoV-2 S pseudotyped viruses

The 2nd generation lentiviral packaging plasmid, psPAX2 (Addgene), a reporter plasmid pUCGFP-Luc (Addgene), and parental or mutant pAbVec-SARS2-S were transfected into HEK293 cells to produce pseudotyped viruses. Briefly, cells plated in 6-well plates the previous day were transfected with three plasmids (1 ug each per well) using Lipofectamine 2000 (Thermo Fisher, Waltham, MA). Following overnight incubation, media was replaced with fresh media containing 5% FBS, and the cells were further incubated for 48 hrs. Supernatants were collected, and cell debris was removed by centrifugation at 400xg for 10 min. Quantitation of pseudotyped viruses was performed using an HIV p24 assay kit (Takara Bio) or ELISA for SARS-CoV-2 S (Sino Biological, Wayne, PA) before storing at -80°C.

### Pseudotyped virus entry assays

To study the entry efficiency of parental or mutant S in cells expressing human or animal ACE2, HEK293 cells or CRFK cells expressing human or animal ACE2 were infected with pseudotyped virus carrying parental or mutant S protein. Briefly, cells plated the previous day were infected with each pseudotyped virus at a multiplicity of infection (MOI) of approximately 1 based on the p24 ELISA for pseudotyped virus preparation. Cell lysates were prepared at 48 h after infection, and firefly luciferase activity was measured on a luminometer (GloMax 20/20, Promega, Madison, WI). Fold change over the parental pseudotyped viruses was calculated for each mutant pseudotyped virus.

### The replication kinetics of SARS-CoV-2 variant strains in cells

SARS-CoV-2 strains USA/WA1/2020 (lineage A), USA/NY-PV08410/2020 (lineage B.1), USA/CA_CDC_5574/2020 (lineage B.1.1.7; alpha variant) and South Africa/KRISP-K005325/2020 (lineage B.1.351; beta variant) were acquired from BEI Resources (Manassas, VA, USA; Supplemental Table 3). Virus stocks were prepared by passaging on either Vero-TMPRSS2 (WA-A, NY-B.1, CA-B.1.1.7, SA-B.1.351) or Calu3 cells (SA-B.1.351), and titers were determined using Vero-TMPRSS2 cells for tissue culture infectious dose 50% per mL (TCID_50_/mL) calculated using the Spearman-Karber method. Virus stocks were sequenced by next generation sequencing (NGS) using the Illumina MiSeq. All variant stock viruses retained the original identified mutations and were in consensus with the original sequenced strains in GenBank. A mutation (PWRAR) in the S furin cleavage site of the SA/KRISP-K005325/2020 Vero-TMPRSS2 passage 1 stock virus was detected as consensus; this stock was used for inoculation of hamsters. The SA/KRISP-K005325/2020 stock was subsequently passaged on Calu3 cells and NGS results showed this stock to contain only 13% of the furin site mutation; this stock was used for the *in vitro* virus replication kinetic experiments.

### SARS-CoV-2 variant replication kinetics in human and hamster ACE2 expressing CRFK cells

SARS-CoV-2 variant replication kinetics were performed with CRFK-human ACE2 or CRFK-hamster ACE2 cells. Cells were inoculated with approximately 0.01 MOI of each virus strain and cell culture supernatants collected at 24, 48 and 72 hours post infection (hpi). Inoculum (defined as 0 hpi) and the time point collected supernatants were then titrated on Vero-TMPRSS2 cells.

### Infection of SARS-CoV-2 variants in the Hamster model

Hamsters (lineage A, *n=8*; B.1.1.7, *n=4*; B.1.351, *n=8*) were inoculated intranasally with 1×10^5^ TCID_50_/mL of virus in 0.1 mL DMEM. Half of the hamsters were humanely euthanized and necropsied at 3 days post challenge (dpc) and the remaining at 5 dpc. Nasal wash was collected from all hamsters at both 3 and 5 dpc and lungs were collected at necropsy. Nasal wash samples were vortexed and stored at -80°C until analysis. Lung homogenates were prepared by homogenizing 200 mg of tissue in a tube containing 1 mL DMEM and a steel bead with a TissueLyser LT (Qiagen, Germantown, MD, USA) for 30 s at 30 hertz, repeated 3 times. Nasal wash and lung homogenates were filtered through a 0.2 μm filter prior to virus titration on Vero E6-TMPRSS2 cells.

### Neutralization assay of convalescent or virus-infected sera from human, cat or rabbit against pseudotyped virus expressing SARS-CoV-2 parental or mutant S protein

The effects of the mutations in the S on antibody neutralizing activity were examined with a panel of SARS-CoV-2 serum samples from humans, cats, and rabbits and pseudotyped virus carrying parental and mutant SARS-CoV-2 S. Serial 2-fold dilutions starting at either 1:12.5, 1:25 or 1:50 of each heat-inactivated serum sample were mixed with a constant amount of each of the pseudotyped viruses carrying parental or mutant S and incubated for 1 h at 37°C. Then, the serum-pseudotyped virus mixture was transduced onto CRFK-human ACE2 cells. Cell lysates were prepared at 48 h after transduction and the relative luminescent units (RLU) were measured. Inhibition curves with each serially diluted serum were generated and the neutralizing titers were calculated as the reciprocal of the serum dilution that showed 50% reduction using GraphPad Prism Software version 6 (San Diego, CA).

### Structural modeling of ACE2-RBD interactions

Protein structural models were produced for the ACE2 interaction with the S protein Receptor Binding Domain (RDB) using the ACE2 sequences for each species listed in Supplemental Table 1 using SWISS-MODEL [90]. The PDB entry 6LZG [91] was used as a template for model building. The ACE2 sequence for each species, corresponding to residues found in the 6LZG template, were used and either the parental SARS-CoV-2 RBD sequence or the mutated sequence (N501Y-K417N-E484K) was added as a hetero target. Structures were analyzed and images created using PyMOL [92].

### Statistical analysis

Statistical analysis was performed using GraphPad Prism Software version 6 (San Diego, CA). One way ANOVA followed by Tukey post-hoc test on the log10 transformed firefly luminescent units, or neutralization titers was used to compare the parental and mutant pseudotyped viruses. To identify significant differences between ACE2-expressing cell cultures or hamsters infected with the different SARS-CoV-2 strains, virus titer data was first log10 transformed and row means and standard deviations were calculated. The data was then analyzed by two-way ANOVA, followed by Tukey’s multiple comparisons test; statistical differences are indicated as * (*p*<0.05). Data are representative of at least two independent experiments. To identify significant differences between groups of hamsters challenged with the different SARS-CoV-2 strains, virus titer data was first log10 transformed and row means and standard deviations were calculated. The data was then analyzed by two-way ANOVA followed by Tukey’s multiple comparisons test and statistical differences indicated as * (*p*<0.05). Data are representative of at least two independent experiments.

## Acknowledgement

The authors thank David George, Emily Gilbert-Esparza, and Yonghai Li for technical assistance, Craig W. Vander Kooi for guidance on structural modeling, and the staff of the Biosecurity Research Institute for assistance with BSL-3 experiments. This research was partially supported the National Institutes of Health R01 award AI109039 (KOC). This investigation also received financial support through grants from the National Bio and Agro-Defense Facility (NBAF) Transition Fund from the State of Kansas (JAR), the AMP Core of the Center of Emerging and Zoonotic Infectious Diseases (CEZID) from National Institute of General Medical Sciences (NIGMS) under award number P20GM130448 (JAR, IM), the NIAID Centers of Excellence for Influenza Research and Surveillance under contract number HHSN 272201400006C (JAR), the World Health Organization (WHO) through a German Federal Ministry of Health (BMG) COVID-19 Research provided to the WHO R&D Blueprint, the NIAID supported Center of Excellence for Influenza Research and Response (CEIRR, contract number 75N93021C00016 to JAR), and the USDA Animal Plant Health Inspection Service’s National Bio- and Agro-defense Facility Scientist Training Program (CM). The authors alone are responsible for the views expressed in this publication and they do not necessarily represent the views, decisions, or policies of the funding agencies.

## Author Contribution

Conceived and designed the experiments: YK, KOC, JAR. Performed the experiments: YK, NNG, DAM, KDP, DB, JDT, IM, CDM. Analyzed the data: YK, KOC, NNG, DAM, JAR. Wrote/revised the paper: YK, NNG, DAM, KOC, JAR.

---

**Supplemental Table 1.**
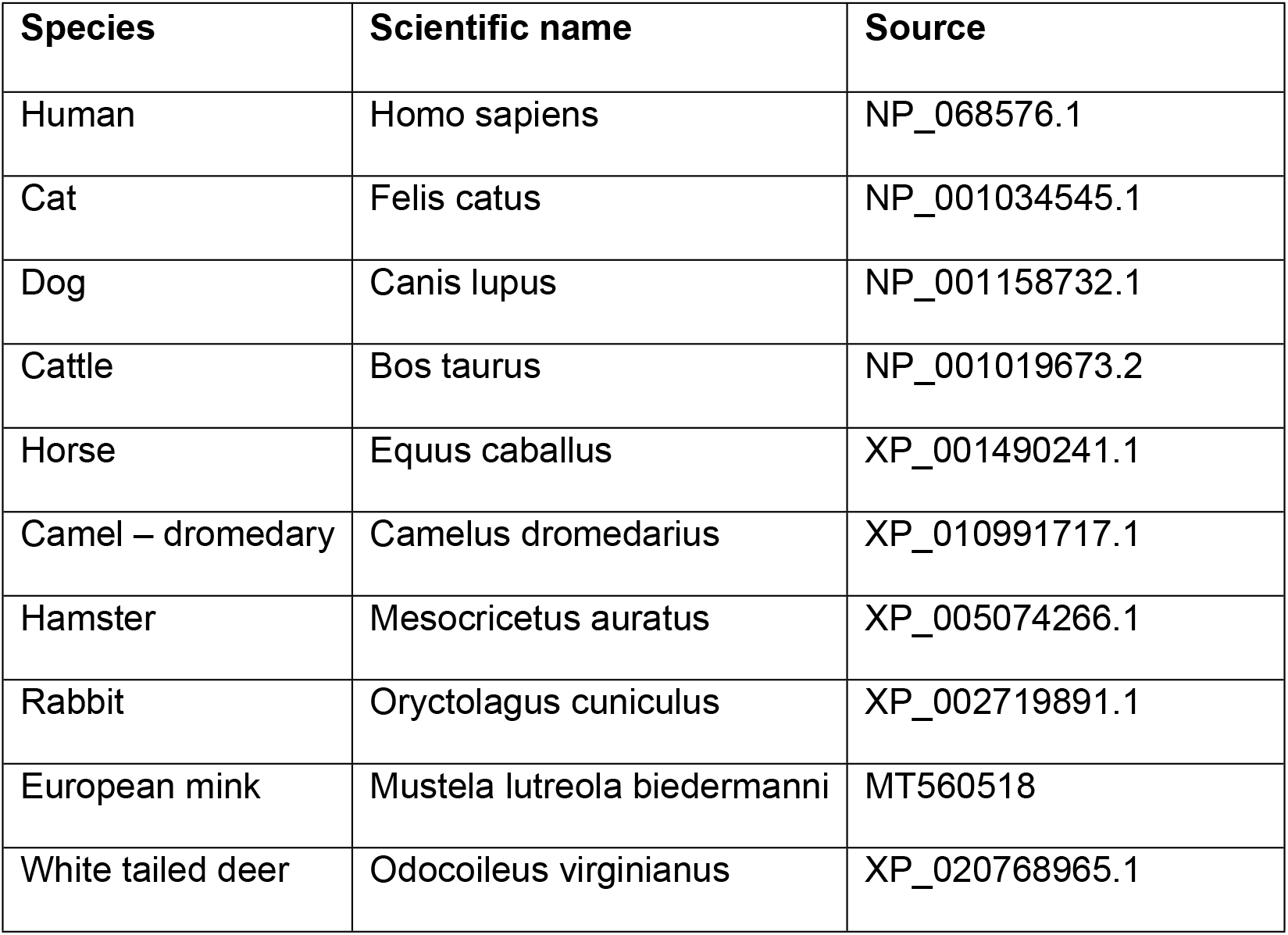
Sources of human and animal ACE2 gene sequences.

**Supplementary Table 2.**
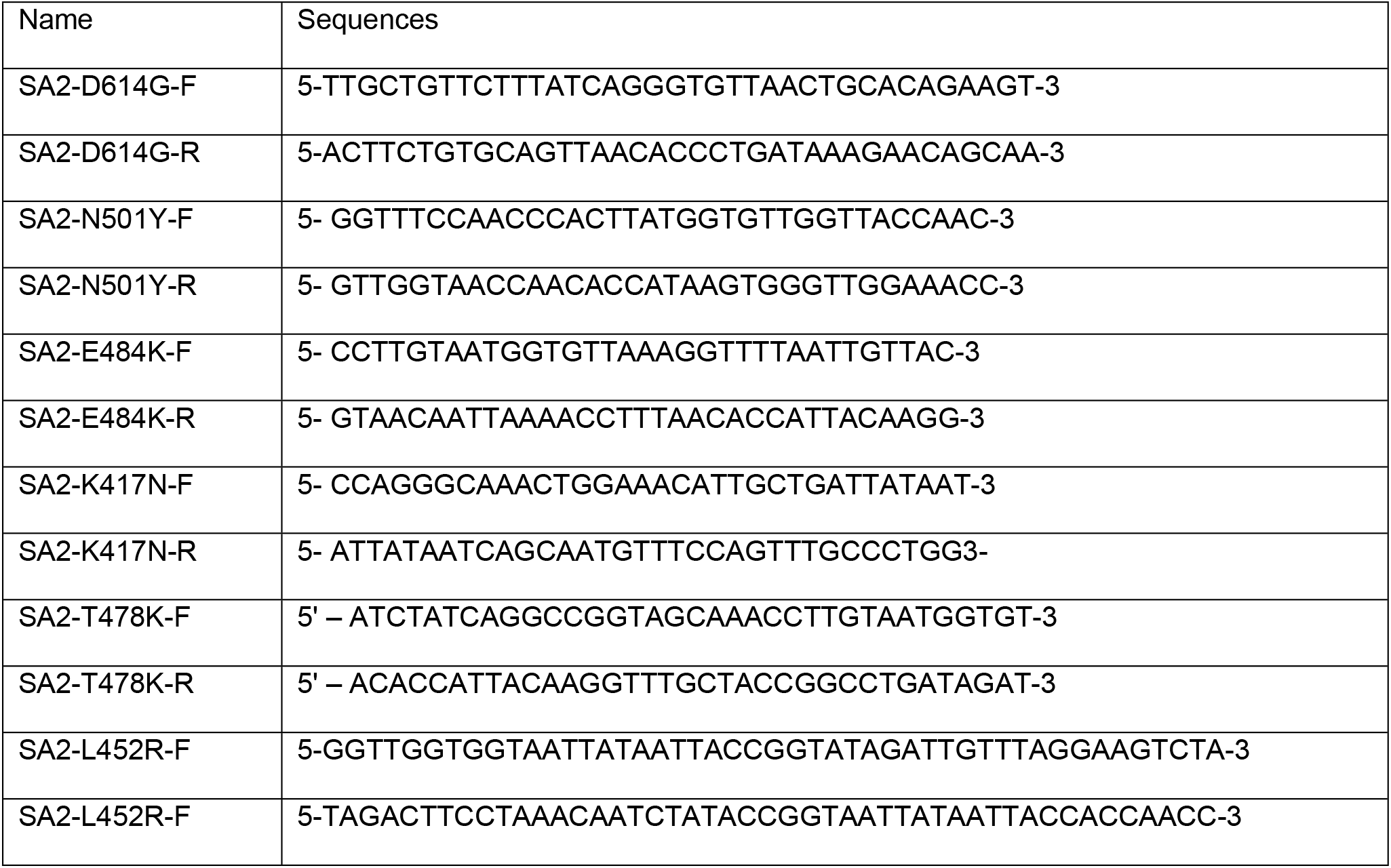
Primers used for mutagenesis of the SARS-CoV-2 S gene.

**Supplementary Table 3.** SARS-CoV-2 strains used for *in vitro* replication kinetics and hamster infection experiments.

| Strain | BEI number | Lineage | Clade | Spike Mutations | Variant |
| --- | --- | --- | --- | --- | --- |
| USA/WA1/2020 | NR-52281 | A | S | na |  |
| USA/NY- PV08410/2020 | NR-53514 | B.1 | GH | D614G | B.1 |
| USA/CA CA_CDC_5574/2020 | NR-54011 | B.1.1.7 | GR | D614G, N501Y | Alpha |
| South Africa/KRISP-K0053252020 | NR-54009 | B.1.351 | GH | D614G, N501Y,<br>E484K, K417N | Beta |

na=not applicable

